# Evolving Bacterial Fitness with an Expanded Genetic Code

**DOI:** 10.1101/169409

**Authors:** Drew S. Tack, Austin C. Cole, R. Shroff, B.R. Morrow, Andrew D. Ellington

**Author notes:** Corresponding Author: Andrew D. Ellington, PhD.

## Abstract

Evolution has for the most part used the canonical 20 amino acids of the natural genetic code to construct proteins. While several theories regarding the evolution of the genetic code have been proposed, experimental exploration of these theories has largely been restricted to phylogenetic and computational modeling. The development of orthogonal translation systems has allowed noncanonical amino acids to be inserted at will into proteins. We have taken advantage of these advances to evolve bacteria to accommodate a 21 amino acid genetic code in which the amber codon ambiguously encodes either 3-nitro-L-tyrosine or stop. Such an ambiguous encoding strategy recapitulates numerous models for genetic code expansion, and we find that evolved lineages first accommodate the unnatural amino acid, and then begin to evolve on a neutral landscape where stop codons begin to appear within genes. The resultant lines represent transitional intermediates on the way to the fixation of a functional 21 amino acid code.

## Introduction

Since the fixation of the genetic code, codon evolution has been confined to the 20 canonical amino acids, with some incursions by selenocysteine and pyrrolysine. Alternative codon tables (e.g. mitochondrial genomes) are likely evolved from the standard codon table and provide evidence that the canonical genetic code can evolve (Knight et al. 2001). A number of theories for the evolution of codon assignment and re-assignment have been proposed (Osawa et al. 1992; Schultz and Yarus 1994; Andersson and Kurland 1998), and the experimental exploration of genetic code malleability has demonstrated the code is not as frozen as once believed, with changes to the code possible over even short evolutionary timeframes (Wong 1983). However, the mechanisms for and intermediary stages of genetic code alteration are largely unknown. Experimental evolution is beginning to address this question, with some recent experiments showing that an expanded set of amino acids can be beneficial (Hammerling et al. 2014). However, a full genomic accounting of the adaptations that occur as a result of alterations to an organism’s standard genetic code has yet to be presented.

Expanding the standard set of proteinogenic amino acids can be accomplished through changes to the underlying translational machinery. Orthogonal translation systems (OTSs) comprising aminoacyl-tRNA synthetase (aaRS)/suppressor tRNA pairs have been developed that do not significantly interact with the host translational machinery or interfere with already occupied portions of the genetic code (Wang and Schultz 2001; Sakamoto et al. 2009; Maranhao and Ellington 2016 Sep 7). Typically, these OTSs allow the incorporation of noncanonical amino acids (ncAAs) by suppressing the amber stop codon (UAG). Cells containing an active OTS exhibit fitness deficits (Wang et al. 2014), likely because any protein terminated by an amber codon can be unnaturally extended. Efforts to knockout the protein responsible for termination at amber codons, release factor 1 (*prfA*), support this claim: some strains lacking *prfA* were found to be viable only when essential genes terminating with amber stop codons were recoded to terminate with the alternative stop codon ochre (Mukai et al. 2010).

This barrier makes the study of genetic code evolution around a new amino acid all but impossible. As work-arounds, evolutionary experiments have been carried out with bacteriophage where the fitness of the organism is irrelevant (Bacher et al. 2003; Hammerling et al. 2014). More extraordinary engineering efforts have led to the development of strains that entirely lack amber codons (Isaacs et al. 2011), which may allow reacquisition of the eliminated codon and insertion of a new amino acid, creating a 21 amino acid genetic code (Hammerling et al. 2016 May 9).

Here we use an alternative approach in which we utilize an engineered β-lactamase (*bla*) that is structurally dependent on OTS incorporation of the ncAA 3-nitro-L-tyrosine (3nY).^9^ This ‘addicted’ *bla* has allowed us to carry out long term evolution experiments without loss of the underlying OTS. Here we demonstrate that our system stably incorporates an ncAA for 2,000 generations of evolution and we for the first time identify the entire complement of genomic mutations in an organism that lead to improved fitness in the presence of an enforced 21 amino acid code.

## Results and Discussion

### Experimental set-up

We wished to examine the long-term adaptation and evolution of *E*. *coli* addicted to a ncAA, 3nY. We assembled an OTS for the incorporation of 3nY comprised of a *Methanocaldococcus jannaschii* tyrosyl-aaRS variant that had previously been engineered to be specific for 3-iodo-L-tyrosine (3iY) (Sakamoto et al. 2009) but was also compatible with 3nY (Tack et al. 2016), and the corresponding *M*. *jannaschii* tyrosyl suppressor tRNA in which the anticodon was complementary to the UAG amber stop codon. This OTS enabled ‘addiction’ via a β-lactamase variant (*bla*_TEM-1.9_) that had been previously selected to be dependent upon 3nY incorporation at amino acid position 162 (Tack et al. 2016). However, since *bla*_TEM-1.B9_ with 3nY already conferred resistance to high levels of ampicillin we further engineered *bla*_TEM-1.B9_ to use ceftazidime (CAZ) as a substrate (Palzkill et al. 1994; Stojanoski et al. 2015). The new β-lactamase, *bla*_Addicted_, conferred moderate resistance to CAZ in a 3nY-dependent manner at antibiotic concentrations commonly used in bacterial cultures (3-10 μg mL^-1^) (see **Figure 1**), and therefore allowed us to both retain and challenge the 3nY incorporation by progressively increasing CAZ concentrations during culture.

**Figure 1.**
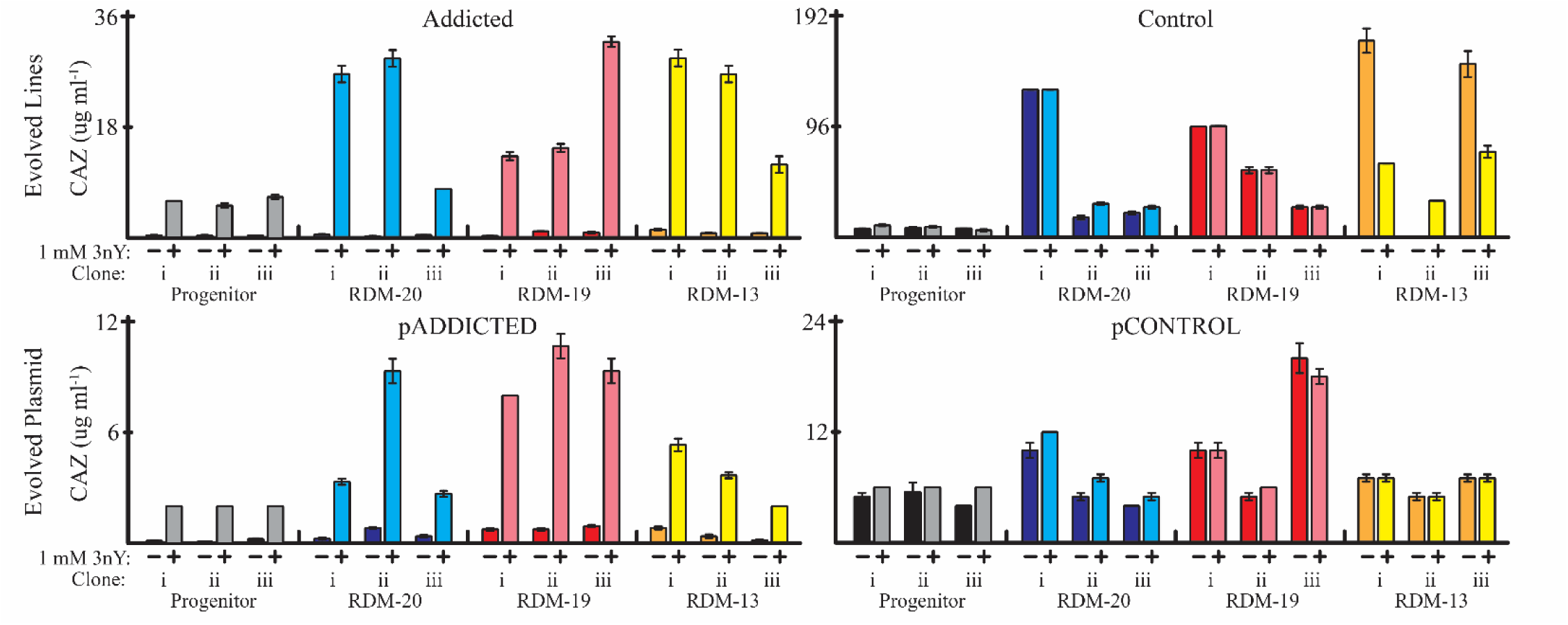
Ceftazidime MICs. MICs of progenitor cells (gray and black) and evolved lines in the absence and presence of 1 mM 3nY. All lines increased MICs during evolution (upper panels, compare gray/black to colored bars). Lines addicted to 3nY remained dependent on 3nY for ceftazidime resistance after 2000 generations (upper left) while control lines never required 3nY for ceftazidime resistance (upper right). Plasmids extracted from evolved lines and transferred to wild-type *E*. *coli* MG1655 showed smaller increases in ceftazidime resistance (lower graphs). Values are the average of biological triplicates, error bars represent s.e.m.

The OTS and *bla*_Addicted_ were assembled together on a plasmid (pADDICTED). We also constructed a control plasmid (pCONTROL) by replacing the 3nY codon (UAG) at position 162 in *bla*_Addicted_ with a phenylalanine codon (UUU), generating *bla*_Control_. Phenylalanine is the only canonical amino acid that produces a functional β-lactamase when replacing 3nY162 in *bla*_TEM-1.B9_(Tack et al. 2016). As expected, pCONTROL conferred CAZ resistance in a 3nY-independent manner (**Figure 1**).

As a chassis for evolution, we chose to use *E*. *coli* strain MG1655 because it is well-characterized, with a sequenced and annotated genome (Blattner et al. 1997). MG1655 is autotrophic for all 20 canonical amino acids allowing for robust growth in amino acid knockout media. MG1655 was transformed with pADDICTED or pCONTROL, and lines were passaged in three different mixtures of amino acids in MOPS-EZ Rich Defined Medium (RDM). The first mixture contained all 20 standard amino acids (RDM-20), the second mixture lacked tyrosine (RDM-19), and the third mixture lacked seven amino acids; serine, leucine, tryptophan, glutamine, tyrosine, lysine, and glutamate (RDM-13) (**Figure 2**). These seven amino acids represent all amino acids encoded by codons accessible through single nucleotide mutations from the UAG stop codon; by limiting the charging of the tRNAs for these amino acids, it should prove more difficult for any single mutation in a codon to be readily suppressed by mutations to tRNA anticodons or by mis-pairing. The RDM-13 media condition also proved a more stringent challenge to growth and evolution. Each media condition was supplemented with 10 mM 3nY, matching the concentration of L-serine, the most abundant amino acid in RDM.

**Figure 2.**
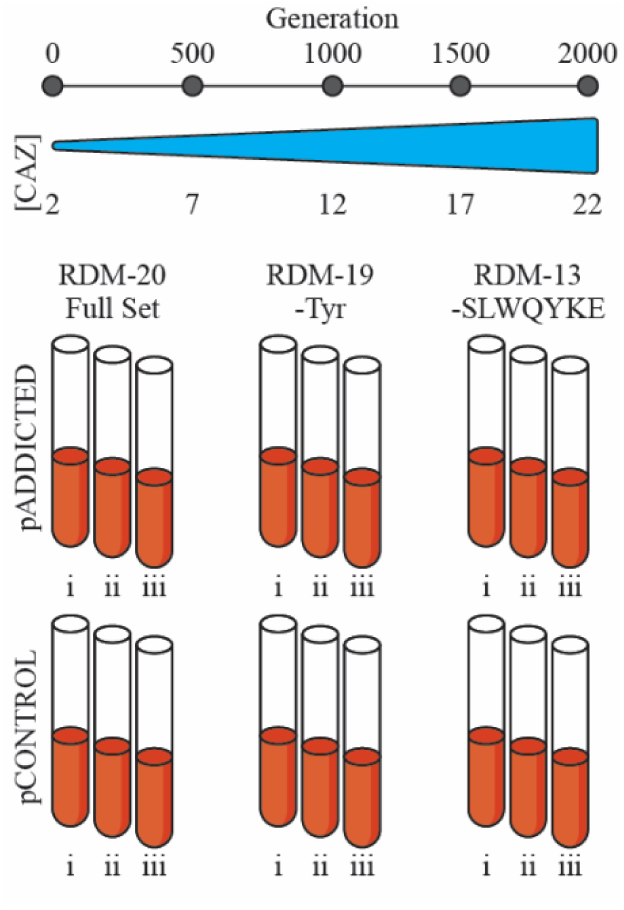
Evolution set up. *E*. *coli* strain MG1655 containing either pADDICTED or pCONTROL was evolved in biological triplicates (i, ii, and iii) for 2000 generations in one of three rich defined media conditions each supplemented with 10 mM 3nY. The first (RDM-20) contained all 20 canonical amino acids, the second (RDM-19) lacked tyrosine, and the third (RDM-13) lacked seven amino acids (serine, leucine, tryptophan, glutamine, tyrosine, lysine, and glutamate). During evolution ceftazidime concentration was increased to provide a fitness burden and enforce OTS activity.

### Experimental evolution of bacteria in the presence of 3nY

We initiated selections with three independent clones containing pADDICTED, and three independent clones containing pCONTROL. These lines were denoted as Addicted(i), Addicted(ii), Addicted(iii), and Control(i), Control(ii), Control(iii) (**Figure 2**). Each clone was passaged in each of the three different amino acid environments described above, and evolved lines are identified by the evolutionary media condition and the progenitor clone from which it was initiated (e.g. Addicted-20(i), Addicted-19(i)…Control-13(iii)).

The cultures were passaged daily by inoculating 5 mL of RDM with 1 μL of overnight growth. This resulted in approximately 12.5 generations per daily passage. While passaging, CAZ concentration was increased at a rate of 1 μg mL^-1^ per 100 generations to a final concentration of 22 μg mL^-1^ to provide evolutionary pressure and ensure enforcement of 3nY dependence. While wild-type MG1655 was capable of growth in all media conditions, the progenitor clones transformed with pADDICTED or pCONTROL proved largely incapable of growth in RDM-13 when supplemented with 3nY, even in the absence of CAZ (**Table 1**, **Supplementary Figure 1**), so for the first 125 generations (10 passages) in RDM-13 the media was supplemented with 25% RDM-19. After these initial 10 passages, all lines were capable of growth in RDM-13.

**Table 1.** Generation rates of *E*. *coli* MG1655 (hr^-1^) in experimental conditions.

| Plasmid | Media | Progenitor |  |  | Evolved |  |  |
| --- | --- | --- | --- | --- | --- | --- | --- |
|  |  | 0 mM | 10 mM 3nY | 10 mM 3iY | 0 mM | 10 mM 3nY | 10 mM 3iY |
| None | RDM-20 | 1.64(.03) | 1.32(.03) | 1.53(.04) |  |  |  |
|  | RDM-19 | 1.46(.07) | 1.25(.01) | 1.46(.07) |  |  |  |
|  | RDM-13 | 1.22(.01) | 0.46(.12) | 1.28(.04) |  |  |  |
| pADDICTED | RDM-20 | 1.65(.05) | 1.42(.04) | 1.50(.03) | 1.77(.08) | 1.69(.05) | 1.67(.04) |
|  | RDM-19 | 1.42(.02) | 1.28(.03) | 1.23(.06) | 1.69(.09) | 1.52(.07) | 1.48(.09) |
|  | RDM-13 | 1.00(.03) | 0.05(.05) | 1.09(.04) | 1.35(.06) | 1.26(.05) | 1.40(.05) |
| pCONTROL | RDM-20 | 1.41(.07) | 1.27(.02) | 1.34(.05) | 1.82(.09) | 1.64(.03) | 1.78(.03) |
|  | RDM-19 | 1.30(.04) | 1.19(.02) | 1.17(.06) | 1.71(.10) | 1.60(.06) | 1.66(.04) |
|  | RDM-13 | 0.94(.05) | 0.14(.09) | 1.14(.03) | 1.40(.04) | 1.38(.05) | 1.41(.05) |

At the conclusion of 160 passages, corresponding to approximately 2000 generations, we selected a single clone from each evolved line and condition and sequenced the bacterial genomes of the single clone as well as the entire bacterial culture using the HiSeq4000 Illumina platform. Selected cells were, in general, genotypically representative of the average bacterial population. We further characterized phenotypic parameters of the selected evolved and progenitor cells.

### Generation rates

We measured the generation rates of progenitor cells in all three media conditions and the generation rates of the evolved lines in the media in which they were evolved. In general, generation rates were measured with and without 3nY, and with and without 2 or 22 μg mL^-1^ CAZ, generation times were calculated based on generation rates in the absence of CAZ and with or without 3nY.

The generation rate of the wild-type MG1655 was slowed by the removal of amino acid supplements (generation rate in RDM-20 compared to RDM-13), as well as by the addition of 3nY to the media (**Table 1**). Cells containing pADDICTED or pCONTROL had similar generation rates to wild-type MG1655 in RDM-20, and RDM-19 irrespective of whether 3nY was added to the media (**Table 1**). However, in RDM-13, the addition of 3nY suppressed generation rates, and the addition of the OTS further exacerbated this fitness challenge; compare, for example, the generation rate of cells without a plasmid in RDM-13 (0.46 generations per hour), with the growth of cells in RDM-13 when containing pADDICTED (0.05 generations per hour) or pCONTROL (0.14 generations per hour). It appears fitness is impacted by amino acid limitations and the addition of the ncAA, and this impact is enhanced if the ncAA can be incorporated. In other words, there are multiple components to cellular fitness in the presence of the ncAA, some of which depend on translation, and some of which do not.

Over the course of evolution, the generation rates of all lines increased in all conditions, except for Control-13(ii), which showed no growth in media lacking 3nY (**Figure 1, Supplementary Figure 1**). The relative fitness effects associated with media conditions remained consistent even after evolution; generation rates were highest in RDM-20 without ncAA, and greater metabolic stresses with decreasing numbers of amino acid supplements resulted in decreases in generation rates. Surprisingly, though, the large growth deficits that were seen with RDM-13 could be largely overcome during evolution.

Since the OTS used is capable of incorporating the ncAA 3iY as well as 3nY, we also tested generation rates of all lines with 3iY. Interestingly, the parent cells were much less affected by growth in 3iY (**Table 1**; compare for example RDM-13 generation rates in the presence of 3nY versus 3iY). This was one of the reasons that 3nY was originally chosen as a evolutionary challenge; it was thought that any mutations that ameliorated the impact of the amino acid on metabolism or translation would be quickly and thoroughly selected. Evolved lines appeared to accommodate 3nY as well as they accommodated 3iY, indicating that whatever specific 3nY toxicities may have been present at the outset have been overcome, and that the cells can now more generally incorporate tyrosine analogues.

### Evolution of antibiotic resistance

During the course of passaging the cultures, CAZ concentrations were increased to levels beyond the MIC of the progenitor cells (the initial MIC of lines was approximately 2-6 μg mL^-1^, while the final CAZ concentration challenge was 22 μg mL^-1^). Bacterial survival at increasingly higher CAZ concentrations indicated that CAZ resistance was evolving in the lines. We measured the MIC of each evolved clone with and without 3nY (**Figure 1**). The addicted lines remained dependent on 3nY for improved ceftazidime resistance, while the MICs of control lines increased in a 3nY-independent manner.

All nine of the addicted cell lines acquired at least a single mutation in *bla*_Addicted_ during evolution (**Table 2**). Several of these mutations are known or expected to be stabilizing mutations, while others are known to expand substrate specificity of *bla*_TEM-1_ (Perilli et al. 2005; Bershtein et al. 2008; Kather et al. 2008; Salverda et al. 2010; Jacquier et al. 2013; Shcherbinin et al. 2017). Other mutations are specific to *bla*_TEM-1.B9_, which originally included a number of substitutions relative to the wild-type *bla*_TEM-1_ that addicted it to 3nY. The substitution T139I in Addicted-20(i) and Control-19(ii), is more similar to the original wild-type residue, leucine, but does not reduce the 3nY dependence of the enzyme. In contrast, only five control lines acquired mutations in *bla*_Control_: Control-20(i), -20(ii), -20(iii), -19(i), and -19(iii). The higher rate of mutation for *bla*_Addicted_ relative to *bla*_Control_ indicates that the enzyme dependent upon the ncAA was not initially as fit as its non-addicted counterpart, especially in the presence of increasing antibiotic concentrations.

**Table 2.** Mutations found in *bla*_Addicted_ and *bla*_Control_ in evolved lines.

| .Mutation | Lines | Effect | Previously Identified |
| --- | --- | --- | --- |
| V33I | Addicted-13(ii) | - | (Salverda et al. 2010; Salverda et al. 2011) |
| Q39K | Control-20(ii) | Expanded Substrate Activity | (Shcherbinin et al. 2017) |
| G92D | Addicted-20(i), -20(ii), -19(ii) | Stabilizing | (Salverda et al. 2010; Figliuzzi et al. 2016) |
| T139I | Addicted-20(i), Control-19(ii) | *bla*<sub>TEM-1.B9</sub> Specific | - |
| T140K | Addicted-19(ii), -19(iii), -13(iii) | - | - |
| M152I | Control-20(i) | *bla*<sub>TEM-1.B9</sub> Specific | - |
| H153R | Addicted-20(ii) | Stabilizing | (Blazquez et al. 2000; Guthrie et al. 2011) |
| M155I | Control-20(iii) | - | (Barlow and Hall 2003; Thai et al. 2009; Salverda et al. 2010) |
| M182T | Addicted-19(i) | Stabilizing | (Kather et al. 2008; Jacquier et al. 2013; Shcherbinin et al. 2017) |
| A184V | Addicted-13(i) | Expanded Substrate Activity | (Forcella et al. 2010; Corvec et al. 2013) |
| T200P | Addicted-20(i) | - | - |
| A224V | Control-19(i) | Stabilizing | (Kather et al. 2008) |

The phenotypic impact of the β-lactamase mutations was further examined by purifying pADDICTED and pCONTROL from the evolved lines and transforming them into wild-type MG1655. These evolved plasmids generally conferred higher MICs than the progenitor plasmids (**Figure 1**). Notably, pADDICTED from lines Addicted-20(ii), -19(i), -19(ii), -19(iii), and -13(i) yielded MIC increases greater than 2-fold the original MIC. These plasmids all contained β-lactamase mutations known or suggested to have stabilizing effects (**Table 2**). However, β-lactamase mutations cannot fully explain MIC changes, as the original lines are generally more resistant to CAZ than are wild-type MG1655 cells with the evolved plasmids transformed (compare upper panels of **Figure 1** with lower panels). Moreover, *bla*_Addicted_ in lines Addicted-19(iii) and Addicted-13(iii) each have only a single, identical mutation (T138K) yet confer different MICs when transformed into wild-type MG1655.

Beyond mutations in the β-lactamase gene itself, increases in MIC may have also been the result of other plasmid or genomic mutations that altered antibiotic resistance through other mechanisms. For example, the plasmid from Addicted-13(iii) has a mutation in *repC*, which can affect copy number and in turn antibiotic resistance (Haring et al. 1985; San Millan et al. 2015). In addition, genomic mutations occurred in genes that are known to have a role in antibiotic tolerance. Four lines (Addicted-20(i), -20(iii), -19(iii), and Control-20(iii)) had mutations in *envZ* (**Table 2**), a histidine kinase that regulates *ompF* and *ompC* expression, which in turn alter membrane porosity and have been tied to β-lactam resistance (Jaffe et al. 1982; Couce et al. 2015). In the same vein, Addicted-13(iii) has a mutation directly in *ompF*. Mutations to *cyaA* or *crp*, two related proteins directly or indirectly involved in transcriptional regulation, occurred in eight evolved lines (Addicted-20(ii), - 19(ii), -19(iii), -13(i), -13(iii) and Control-19(i), -19(iii), -13(iii); inactivation of these genes has been shown to produce resistance to β-lactams (Jaffé et al. 1983; Ruiz and Levy 2011). The *opgG* gene was mutated in Addicted-20(iii), -19(i), and Control-20(ii), and is involved in conferring resistance to antibiotics (Hanoulle et al. 2004). Finally, lipopolysaccharide (LPS) expression has been tied ceftazidime treatments, especially at or near MIC levels, and several lines had mutations in LPS biosynthesis and maintenance genes. Notably, seven lines had IS insertions in the *waa* operon that encodes the core oligosaccharide of LPS, including *waaO* (Addicted-20(ii), -19(i), and Control-20(ii), -20(iii)), *waaQ* (Control-20(i)), *waaP* (Addicted-13(i)), and *waaB* (Control-19(ii)). Also, three lines had mutations in the LPS-related gene *galU*, (Control-19(i), -13(i), -13(iii)), and one line had a mutation in *lptD*, which is involved in the assembly of LPS at the surface of the cell (Control-13(ii)) (Pagani et al. 1990; Leying et al. 1992).

Beyond the more frequent stabilization of *bla*_Addicted_, there were few sequence substitutions that were specific to lines addicted to 3nY relative to controls that were not addicted to 3nY. None of the lines appear to use an expanded genetic code to increase fitness through ceftazidime resistance.

### Retention of the OTS

One of the primary questions we were addressing was whether lines that were addicted to the OTS would retain and utilize the ncAA differently or better than lines that were not addicted to the OTS. The 3nY dependent MICs (**Figure 1**) strongly suggests that a functional OTS has been retained in all addicted lines. To directly evaluate the functionality of the OTS we measured UAG suppression efficiency with a GFP reporter system that contained a tyrosine (TAT), amber (TAG), or ochre (TAA) codon at position Y39 of GFPmut2 (Cormack et al. 1996). We tested the each GFP reporter in progenitor cells as well as each of the evolved lines (**Supplementary Figure 2**). All of the addicted lines remained capable of efficient amber suppression (>10% 3nY incorporation at UAG codons) (**Figure 3**). In contrast, three of the nine control cell lines (Control-20(i), -19(ii), -19(iii)) had lost UAG suppression capability (<1% 3nY incorporation) (**Figure 3**, see ‘∗’). Surprisingly, UAG suppression efficiency generally dropped in both addicted and control lines relative to their progenitor cells.

**Figure 3.**
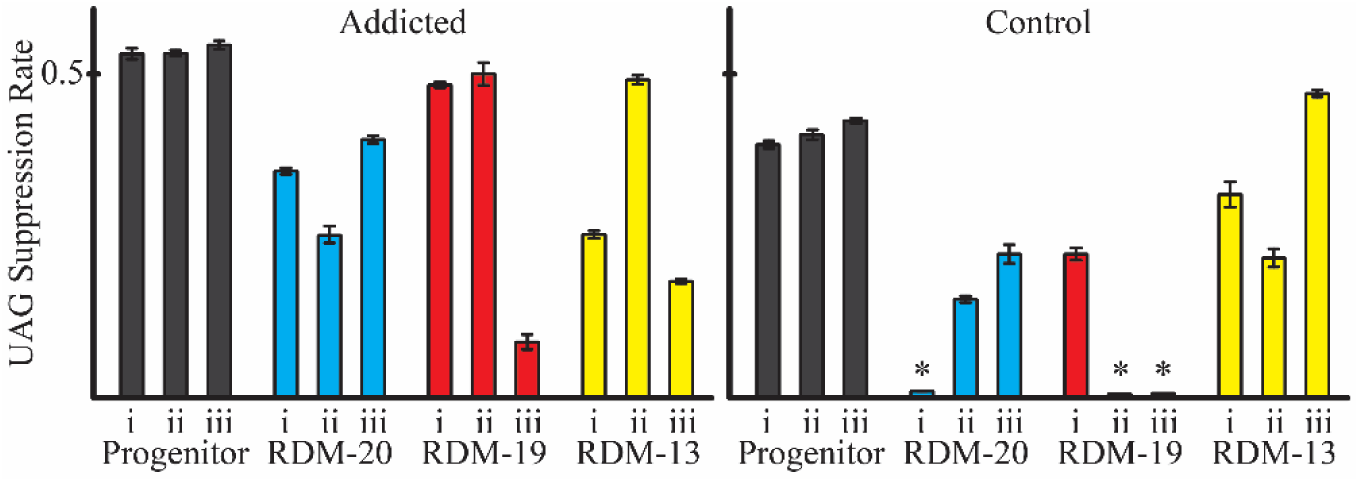
UAG Suppression Efficiency. OTS activity of progenitor cells (black) and evolved lines was determined by measuring UAG suppression efficiency. All addicted lines maintained an active OTS (left), while three control lines lost OTS activity over 2000 generations (see ‘^∗^’). Values are the average of four biological replicates, error bars represent s.e.m.

Of the addicted lines, only Addicted-19(i) acquired mutations in the OTS, acquiring a T->G mutation in the tRNA processing machinery at position T(-28). In contrast, OTS mutations occurred in Control-20(i), -19(i), - 19(ii), -19(iii), three of which had lost OTS activity. Of these, lines Control-20(i) and Control-19(ii) have aaRS mutations: Control-20(i) has a frameshift in the aaRS resulting in a UGA ochre stop codon at amino acid position 123, which leads to the complete loss of UAG suppression, and Control-19(ii) contains an IS-mediated insertion in the aaRS, a mechanism that has previously been observed to inactive OTS machinery (Wang et al. 2014). Control-19(i) and Control-19(iii) each had mutations in the tRNA. The tRNA of the OTS from Control-19(i) contained a G26C substitution in the hinge between D-stem and C-stem (**Figure 4**, yellow), while OTS tRNA from Control-19(iii) has a G42A substitution in the C-stem of the tRNA (**Figure 4**, red). The latter mutation is predicted to compromise tRNA structure, as determined using mfold (Zuker 2003) and tRNA prediction software (Schattner et al. 2005) (**Figure 4**). This tRNA mutation likely explains the complete loss of UAG suppression in Control-19(iii) (**Figure 3**).

**Figure 4.**
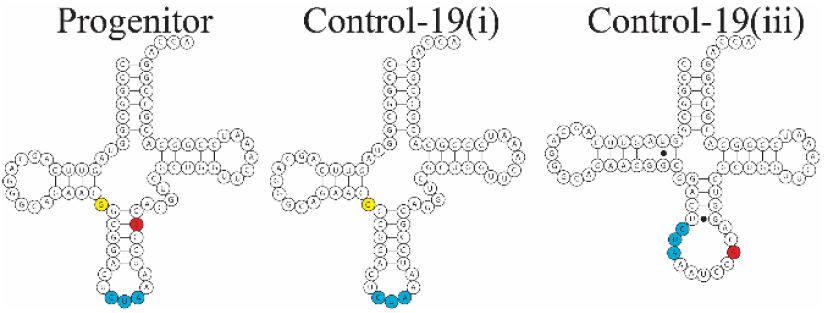
Predicted tRNA structures. Mutations to the OTS tRNA occurred in two lines during evolution. Computationally predicted tRNA structures from these evolved lines shows substitution G26C (yellow) found in Control-19(i) (center) is not likely to affect general structure or the anticodon (blue), while substitution G42A (red) in line Control-19(iii) is predicted to disrupt tRNA structure (right).

Overall, either suppression in general or the specific incorporation of 3nY puts cells at a fitness disadvantage, and in the absence of addiction to the ncAA this trait can be readily lost. The evidence suggests that suppression in general is less toxic than 3nY incorporation. First, the parental strain can take up and incorporate 3iY into proteins without a large diminution of generation rate (**Table 1**, compare generation rates in the presence of 3iY with no plasmid, and with pADDICTED or pCONTROL, each carrying an active OTS for 3iY incorporation), while 3nY is much more deleterious to growth. Second, in the absence of addiction the OTS is readily lost or compromised during evolution (**Figure 3**, compare suppression efficiencies in ‘Addicted’ and ‘Control’ panels). Additionally, previous work also supports the hypothesis that suppression is not particularly toxic, given the existence of natural amber suppressors which have minimal effect on fitness or the transcriptome (Herring and Blattner 2004).

### Genomic adaptations

As anticipated, the number of mutations acquired by a line scaled with the stringency of selection. Full genome sequencing and analysis revealed that addicted lines acquired more ORF-affecting mutations than controls (**Table 3**); forty-nine ORF-affecting mutations in addicted lines, and forty in the control lines. Additionally, mutations generally increased with decreasing amino acid supplements, with the addicted and control lines together accumulating twenty-two mutations in RDM-20, thirty-one mutations in RDM-19, and thirty-six mutations in RDM-13. Several trends were seen among evolution conditions and the genes affected during evolution.

**Table 3.**
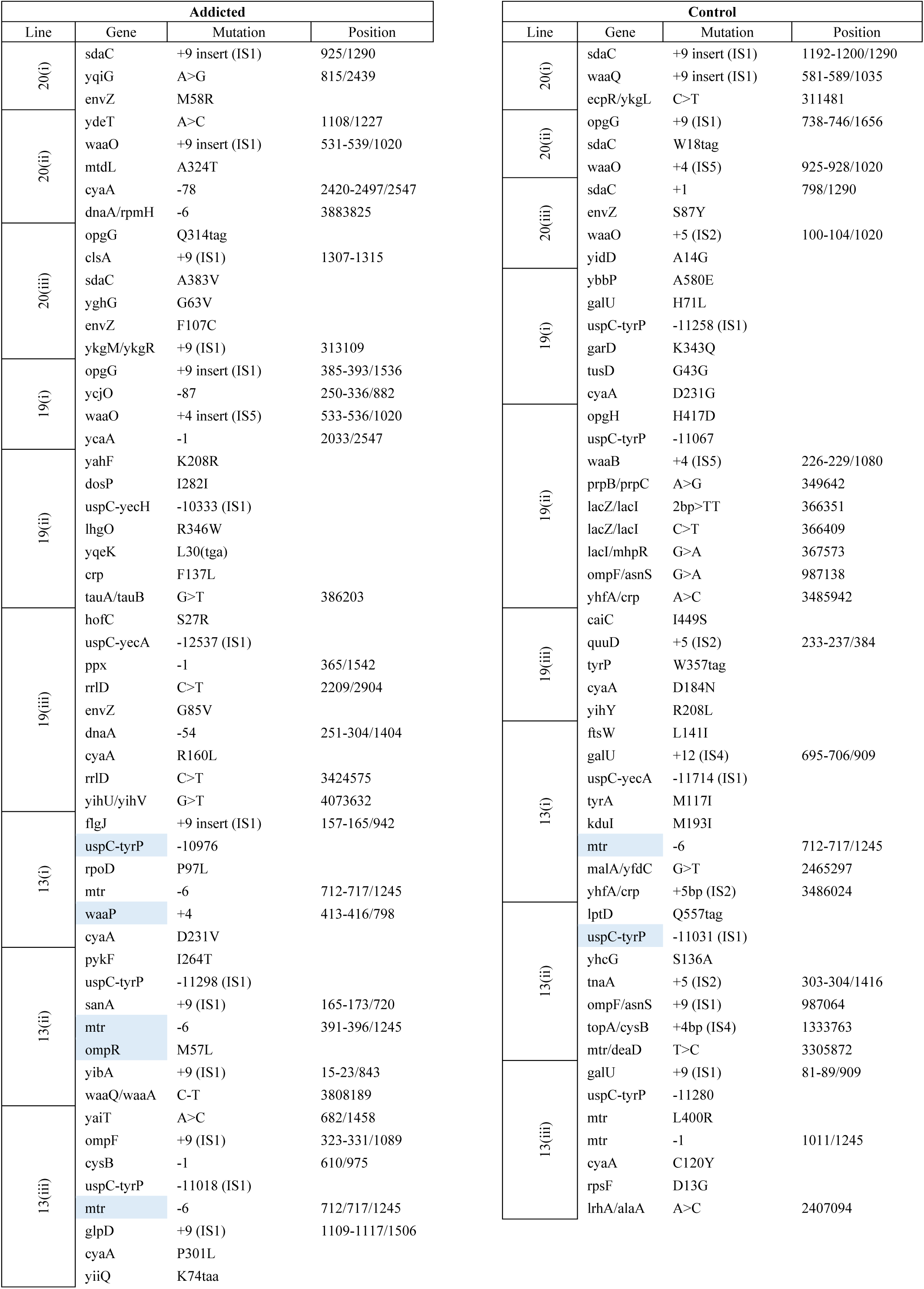
Genomic Mutations of Evolved Lines. Mutations which appeared after 125 generations in RDM-13 are highlighted blue.

Strikingly, the most commonly mutated or deleted ORFs were amino acid transporters in the hydroxyl and aromatic amino acid permease (HAAAP) family. The most commonly affected HAAAP protein was *tyrP*, a tyrosine-specific permease (Wookey and Pittard 1988; Andrews et al. 1991) that was inactivated or deleted in 10 of the evolved lines, including all six evolved in RDM-13, and four of the six lines evolved in RDM-19 (all three control lines, as well as Addicted-19(iii). The most common mechanism (9 of 10 cultures) used to inactivate *tyrP* was an IS1 mediated deletion of a large genomic fragment that excised twelve to fourteen genes (also discussed below), including a portion or the entirety of *tyrP*. In addition, we observed one example of an in-frame stop codon at position W357 of Control-19(iii). The second most commonly mutated HAAAP protein was *mtr*, a tryptophan-specific permease (Brown 1970). The *mtr* mutations were exclusively found in lines evolved in RDM-13, with five of the six RDM-13 lines (all except Control-19(iii)) having a mutated *mtr* gene, and the final having an intergenic mutation in close proximity to *mtr*. Four of the lines acquired in-frame, hexa-nucleotide deletions, which occurred in transmembrane domains (Sarsero and Pittard 1995). Three lines (Addicted-13(i), -13(iii) and Control-13(i)) deleted the same six nucleotides, positions 711-716, excising amino acids A238 and L239 of transmembrane span VII, the fourth line deleted nucleotides 391-396, excising amino acids G131 and F132 of transmembrane span IV. The fifth mutant *mtr* gene contained a single nucleotide deletion at nucleotide position 1011 of *mtr*, resulting in a frameshift from amino acid position 338 onward. Unlike *tyrP*, the *mtr* gene did not mutate in lines grown in tryptophan rich media (RDM-20 and RDM-19). The third most common HAAAP protein that was *sdaC*, a serine specific transporter, which was inactivated in four of the six lines evolved in RDM-20; a mutation of unknown consequence appeared in a fifth line (A384V), while *sdaC* remained fully unaffected in all RDM-19 and RDM-13 evolved lines.

We hypothesized that deactivation of HAAAP transporters may have played a role in reducing the metabolic impact of 3nY, most drastically seen in RDM-13. Evidence for the importance of these mutations for adaptation to growth in 3nY comes from an examination of the mutations that arose in the RDM-13 lines between generations 0 and 125, especially the lines containing the pADDICTED gene, which were initially incapable of growth in RDM-13 and had to be supplemented over their first 125 generations (10 passages) with 25% RDM-19. We sequenced the genomes of the six RDM-13 lines at generation 125 to determine which mutations had occurred early in experimental evolution, and found that five of the six populations had mutations in *mtr* or the *mtr* promoter. These mutations were fixed in three of the five lines, and for the remaining two 22.8% of the Control-13(i) population had a *mtr* mutation, and 44.2% of the Addicted-19(iii) had mutations in *mtr*; both of these mutations were largely fixed by generation 2000. Additionally, the *uspC*-*tyrP* region was excised early from the genomes of Addicted-13(i) and Control-13(ii). Taken together, this data supports the hypothesis that HAAAP deactivation ameliorated 3nY toxicity.

To determine the gross fitness impacts of amino acid transporter deletions we examined the generation rates of *tyrP* and *mtr* knockouts from the Keio collection (Baba et al. 2006) in RDM with and without 3nY. Strains from this knockout collection are based on *E*. *coli* strain BW25113, which is genetically similar to our host strain MG1655, in that both are K-12 isolates that have been cured of the F plasmid and phage lambda (Bachmann 1972; Baba et al. 2006). The *tyrP* and *mtr* knockouts did not affect growth compared to wild-type *E*. *coli* in RDM-20 and RDM-19, but significantly delayed growth in RDM-13, both with and without 3nY (**Supplementary Figure 3**). These results suggest that alterations in amino acid transporter function may have mediated a subtle fitness component, and would likely have been under selection over the multiple generations of evolution that were carried out.

A large (~10 kb) IS1-mediated deletion of the genome occurred in 10 of the 18 evolved lines, including all six RDM-13 evolved lines, as well as four of the RDM-19 evolved lines (Addicted-19(ii), -19(iii) and Control-19(i), -19(ii). This was the most common mechanism used to inactivate the *tyrP* transporter. While no two deletions were identical, the deleted region in general contained: the universal stress protein C gene (*uspC*), which is involved in UV tolerance and the transition to stationary phase during growth in rich media (Gustavsson et al. 2002; Siegele 2005); the trehalose-6-phosphate synthase (*otsA*) and trehalose-6-phosphate phosphatase (*otsB*) genes, which are involved in osmoregulation, and contribute to antibiotic tolerance and oxidative stress prevention (Kuczyńska-Wiśnik et al. 2015); the arabinose transporter subunit genes *(araH*, *araG*, and *araF*), which are responsible for the cross-membrane transport of the sugar L-arabinose (Kolodrubetz and Schleif 1981); and the ferritin iron-storage complex (*ftnB*) and related protein (*ftnA*) genes that are involved, or predicted to be involved, in iron storage (Hudson et al. 1993); and the small membrane protein (*azuC*) that has been identified as a stress response protein (Hemm et al. 2010; Fontaine et al. 2011). In addition, the deletion encompassed the genes for the predicted proteins *yecJ*, *yecH* and *yecR* of unknown function, but potentially playing a role in cell envelope or flagellar formation (Stafford et al. 2005; Paradis-Bleau et al. 2014). Additionally, *yecA* has predicted metal binding properties (Serres et al. 2001). All of these genes were deleted in all ten lines with the exception of *yecA* and *tyrP*, which remained fully intact in Addicted-19(ii).

A final, commonly mutated ORF amongst all lines was adenylate cyclase (*cyaA*), which biosynthesizes cyclic-AMP (cAMP) which is in turn a secondary messenger that activates or inhibits the expression of over 400 genes in *E*. *coli* via the transcriptional regulator *crp* (Khankal et al. 2009). Eight lines evolved mutations in *cyaA* or *crp*, of which seven acquired SNPs resulting in a single amino acid substitution, each unique. The final mutation, in Addicted-20(ii), was a 78 base-pair deletion in the C-terminus of the protein, cleanly deleting the regulatory domain of *cyaA* while leaving the enzymatic domain intact (Roy et al. 1983). Mutations to *cyaA* or *crp* have been shown to impact diverse phenotypes in carbon utilization, stress response, and amino acid metabolism (Gosset et al. 2004), and the acquired mutations in our evolved lines may have ameliorated the fitness effects of ncAA incorporation. As examples of potentially relevant metabolic impacts, when *cyaA* or *crp* were deleted from the genome of *E*. *coli*, the bacteria became incapable of using amino acids as a carbon source and depended on the presence of glucose for growth (Donovan et al. 2013). Additionally, down-regulating or inactivating *cyaA/crp* activity resulted in the constitutive expression of the regulatory proteins sigma E and *Cpx*, which in turn up-regulated chaperone proteins and proteases involved in stress responses, including temperature sensitivity and oxidative stress (Strozen et al. 2005; Donovan et al. 2013).

### Intergenic and noncoding mutations

While the majority of mutations in the evolved lines occurred in protein-encoding sequences, a total of 19 mutations occurred in intergenic and noncoding regions throughout the eighteen evolved lines (**Table 3**). Addicted lines acquired six noncoding mutations, while control lines acquired a total of 13 noncoding mutations, with six occurring in the Control-19(ii) alone. Many of these mutations occurred near genes that also had mutations in their coding regions. For example, while five of the six lines evolved in RDM-13 had likely inactivating mutations in the *mtr* tryptophan transporter gene, the sixth (Control-19(ii)) appears to downregulate *mtr* expression via a SNP that arose in the *mtr* promoter that disrupts a critical position in the Pribnow box (Harley and Reynolds 1987). The *cysB* transcriptional regulator of sulfur metabolism had a frameshift mutation in Addicted-13(iii), and has a IS4-mediated 4-basepair insertion in the promoter of Control-13(ii). Similarly, *ompF* was likely inactivated in Addicted-13(iii) via a polynucleotide insertion, but expression of *ompF* was affected in Control-19(ii) by mutating upstream regulatory sites for *ompR* and *crp* binding, and in Control-13(iii) by a destabilizing IS1-mediated insertion of nine nucleotides in a 5’ hairpin in the mRNA transcript of *ompF* (Papenfort et al. 2006; Podkaminski and Vogel 2010; Gogol et al. 2011). Similarly, *dnaA* gene had a 54-basepair deletion in Addicted-19(iii), and also accrued a mutation of unknown effect 6 bp upstream of the ORF in Addicted-20(ii). Finally, two lines had mutations in the promoter or regulatory regions of *crp*; Control-19(i) had a 5 base-pair insert at a fis-transcriptional regulator site (González-Gil et al. 1998), and in line Control-19(iii) a SNP occurs upstream of *crp*.

### Sampling of new codes

The clone chosen for phenotypic characterization and individual genome sequencing from one control line was conspicuously different than the clones from all other evolved lines. When we assayed the ceftazidime MIC for Control-13(ii) there was minimal to no growth on plates without 3nY, but appropriate growth on 3nY supplemented plates. Later, multiple replicates consistently resulted in a no growth phenotype for line Control-13(ii) in the absence of 3nY during amber suppression assays. Sequencing of this genome revealed an in-frame UAG codon in *lptD*, a Q557(UAG) substitution. *lptD* is an essential gene involved in LPS biosynthesis (Braun and Silhavy 2002; Baba et al. 2006). The apparent essentiality of this mutation suggests that Control-13(ii) has adopted a 21 amino acid genetic code, and this occurred even in the absence of an addicted β-lactamase variant. Comparison to the full culture sequencing revealed this mutation was not representative of the general populace. To our knowledge, this is only the third time an alternative genetic code has evolved in the context of directed evolution, following on the early directed evolution of a *B*. *subtilis* strain that could utilize 4-fluorotryptophan in place of tryptophan, and the more recent directed evolution of bacteriophage T7 in an *E*. *coli* strain that could incorporate 3iY across from amber stop codons.

## Conclusions

Previously, two different approaches have been used to generate organisms with altered or expanded amino acid genetic codes a bottom-up approach where components of the organism were engineered to function with a ncAA, or a top-down approach, where an organism was allowed to evolve in the presence of a ncAA (Bacher et al. 2004). Advances in genome editing and protein structural modeling have made the bottom-up approach feasible, with two recent reports of *E*. *coli* successfully engineered to depend on a ncAA for survival (Mandell et al. 2015; Rovner et al. 2015). It should now be possible to make fully synthetic genomes (Gibson et al. 2010) that are designed to have a chemical dependence on an ncAA throughout the genome, although the ‘amberless’ *E*. *coli* (Lajoie et al. 2013) introduced a number of unanticipated mutations that diminished fitness. Recently, experimental evolution of ‘amberless’ *E*. *coli* without an OTS allowed the ‘amberless’ strain to correct these fitness deficits (Wannier et al. 2017 Jul 12). Alternatively, top-down approaches have previously been used to generate organisms that preferentially function with a ncAA substituting for tryptophan (Wong 1983). Using mutagens and selective growth conditions, *Bacillus subtilis* became dependent on a normally toxic tryptophan analog, 4-fluorotryptophan, with only 106 genomic mutations required to change amino acid preference (Yu et al. 2014). In contrast, an *E*. *coli* tryptophan auxotroph evolved in the presence of 4-fluorotryptophan in place of tryptophan (Bacher and Ellington 2001) could tolerate high levels of the ncAA, but in the end still required tryptophan for growth. A more recent report shows that the bacteriophage T7 will adopt ncAAs into its genetic code to reach new fitness peaks (Hammerling et al. 2014).

The work described herein can be considered a hybrid approach, resting between bottom-up and top-down. We used a single engineered addiction element, *bla*_Addicted_, to preserve an active OTS over evolutionary timeframes within an otherwise unmodified organism. Thus during 2000 generations of evolution bacteria were required to retain 3nY incorporation as part of its genetic code and could fully explore the potential utility of 3nY throughout their proteomes. Overall, our method provides one of the first real world examinations of how a new genetic code is adopted by an organism.

The ability to encode 3nY ambiguously across from stop codons was clearly initially a fitness disadvantage for cells. Several lines of evidence support this conclusion. First, in the absence of the addiction module the OTS was inactivated during evolution. Second, amino acid transporters were frequently inactivated, especially during the initial evolution of cells under the most stringent selection conditions (RDM-13). These mutations likely decreased the uptake of 3nY, and reduced the ambiguity of encoding. Third, bacteria began ameliorating the fitness burden by introducing systemic changes to metabolism that likely improved stress response (mutations in *cyaA*, *crp*, and *envZ*).

Nonetheless, while bacterial fitness and addiction were pitted against one another, addiction proved to be the less malleable of the two, and in the presence of the *bla*_Addicted_ no OTS loss occurred. Indeed, many of the observed sequence changes focused on improving, rather than eliminating or breaking, the addicted β-lactamase. This was not a foregone conclusion: in order to re-establish the canonical 20 amino acid code, evolved lines could have, for example, augmented the rates of ncAA incorporation through modifications to release factor 1 that led to increased termination efficiency or decreased OTS activity while increasing *bla*_Addiction_ expression.. Similarly, even though 326 genomic amber codons were a single mutational step from stop codons that would not be suppressed by the OTS, in over 2000 generations addicted populations fixed no mutations that led away from the 21 amino acid code, despite previous evidence that recoding amber codons to alternative stop codons reduces the fitness effects of obligate UAG suppression.^5,7^(Hammerling et al. 2014)

Our hybrid approach resulted in mutations that led to the accommodation of the ncAA, and thus to a fitness landscape in which in-frame UAG codons could be neutrally adopted, and where those UAG codons could apparently encode either 3nY or 3iY. In many ways, these results with whole organism evolution resemble numerous results from the field of directed protein evolution. Enzymes evolved to change their substrate specificities frequently go from being specialists (working on the wild-type substrate) to generalists (able to accommodate many different substrates), and then finally become a new specialist (preferentially utilizing the new substrate) (O’Brien and Herschlag 1999). In our case, the evolution of the genetic code has preceded from being specialized for 20 amino acids, to being able to accommodate an ambiguous code of 21 amino acids, where amber codons can code for 3nY, 3iY or stop.

Overall, our efforts can thus be taken as molding a strain that is at least neutral to an ambiguous 21 amino acid genetic code (fitness returned to wild-type levels after evolution), and is now more fully prepared for the specification of a new code. As stop codons neutrally accumulate, they eventually invade essential positions in essential genes, such as the *lptD* gene in Control-13(ii) reported here, and the holin gene in previous experiments with T7 phage (Bacher and Ellington 2001; Hammerling et al. 2014). Such neutral invasion ultimately more firmly fixes the new code, and as evolution continues will presumably lead to fixation of a particular 21 amino acid code. This hypothesis is buttressed by the existence of an isolated clone that utilized the full 21 amino acid code in a critical protein, which required suppression for growth.

## Methods

- Evolutionary set up
  - Addiction operon Plasmid backbone from pMMB67EH (Fürste et al. 1986) was amplified using DNA oligos (Integrated DNA Technologies) DT01 and DT02 (**Supplementary Table 1**). The *M*. *jannaschii* OTS and *bla*_TEM-1.b9_ were amplified from a plasmid described previously (Tack et al. 2016) using oligos DT03 and DT04. To convert the penicillinase *bla*_TEM-1.B9_ to the cephalosporinase *bla*_Addicted_, residues 165-167 were converted from WEP to YYG using oligo DT05 with either DT06 or DT07 for *bla*_Addicted_ or *bla*_control_ respectively. Reaction mixtures were transformed into *E*. *coli* TOP10 and selected on LB-agar with 2 μg mL^-1^ CAZ for *bla*_control_, and the same conditions with 10 mM 3nY for *bla*_Addicted_. Samples were sequence verified at University of Texas core facilities using Sanger sequencing. One properly sequenced plasmid of each *bla*_control_ and *bla*_Addicted_ were used as pCONTROL and pADDICTED respectively, and transformed into *E*. *coli* MG1655. Three colonies from each were selected as clones i, ii, and iii for passaging.
  - Media MOPS-EZ Rich Defined Media (RDM, TEKnova) with the full complement of amino acids, as well as the knockout medias (RDM-19 and RDM-13), were prepared according the manufacturers specification. For media preparation, 3nY (Sigma-Aldrich) was added to water immediately after autoclaving for sterility, to a final concentration of 17.24 mM. Once cooled, the 3nY supplemented water was used to complete RDM, and the entire preparation was filter sterilized with Nalgene Rapid-Flow SFCA filtration units. Media was stored at 4°C before use, and moved to room temperature approximately 16 hours before use.
  - Passaging Selected colonies i, ii, and iii, from each transformation were picked and grown in RDM-20 supplemented with 10 mM 3nY and 2 μg mL^-1^ CAZ for 16 hours. Each culture was then used to inoculate three subcultures of RDM-20, RMD-19, and RDM13. Initially, cultures were incapable of growth in RDM-13, but were capable of growth when 25% of media was replaced with RDM-19. This growth condition was used for the first 125 generations, after which cultures were capable of growth in RDM-13. Cultures were passaged every 16-24 hours by transferring 1 μL of previous growth culture into 5 mL of RDM, and grown shaking at 37°C. A 500 μL samples from each line was preserved in 25% glycerol at -80°C at generation 0, 125, 250, and every 250 generations for the duration of evolution.
- Genome Sequencing and assembly After the 2000 generations, 1 μL from each line was streaked onto RDM-agar supplemented with 10 mM 3nY. Two single colonies from each line were selected and grown in 3 mL RDM-20 with 10 mM 3nY with 22 μg mL^-1^ CAZ. Simultaneously, samples from the progenitor cells were grown in similar conditions using 2 μg mL^-1^ CAZ. After overnight growth, glycerol stocks were made of monoclonal cultures, and genomic DNA was isolated from one of the two cultures using bacteria genome miniprep kit (Sigma-Aldrich). Bacterial genomic DNA samples were prepped and sequenced with 150 bp paired ends using the HiSeq 4000 platform, achieving greater than 100x coverage across all samples. Raw reads were processed through trimming and adapter removal using trimmomatic (v0.32) (Bolger et al. 2014 Apr 1). Alignment of sequencing reads and variant calling was performed through the breseq workflow (v0.27.2) (Deatherage and Barrick 2014).
- Growth Curves Glycerol stocks were used to inoculate 2 mL of appropriate RDM and incubated overnight while shaking at 37°C. Then, 1 μL from overnight cultures were used to inoculate 100 μL of media with appropriate concentrations of CAZ and 3nY. A Tecan Infinite M200 pro microplate reader set to 37°C monitored absorbance at 600 nm for approximately 24 hours. Data for each well were trimmed from 0.1 to 0.2 OD after which the rate of logarithmic change in OD was calculated for each line to determine generation rates.
- MICs The six progenitor cells and the eighteen clones from the evolved lines were grown overnight in 3nY enriched MOPS-EZ growth media supplemented with 2 μg mL^-1^ or 22 μg mL^-1^ of CAZ respectively. Overnight cultures were used to inoculate fresh RDM lacking 3nY and without CAZ, and were grown for five hours at 37°C while shaking. Aliquots of 25 μL from each line (corresponding to ~10^7^ cfu) were plated on LB-agar, with or without 1 mM 3nY, in triplicate. Plates were allowed 5 minutes to dry in a sterile biosafety cabinet, and ceftazidime E-test MIC strips (Biomerieux) were added to each plate. Plates were incubated 16 hours at 37°C. MIC values are reported as the lowest concentration on E-test strip at which no bacterial growth occurred.
- GFP Assays The codon for Tyr39 (TAT) in GFPmut2, under control of the *tacI* promoter with *Lac* operator, was changed to the amber codon (TAG), or ochre nonsense codon (TAA) using Gibson cloning with oligo DT10 paired with DT08 (for TAG codon) or DT09 (for TAA codon). Gibson reaction was electroporated into TOP10 *E*. *coli*, and plated on LB-agar supplemented with 50 μg mL^-1^ kanamycin (KAN). Sequences were verified using Sanger sequencing at the University of Texas Core Facility. Glycerol stocks of the eighteen evolved clones and six progenitor clones were used to inoculate 2 mL of LB supplemented with 10 mM 3nY and 2 or 22 μg mL^-1^ ceftazidime (progenitor and evolved cells, respectively), and grown to saturation overnight. A 5 μL aliquot from each was used to inoculate 5 mL of LB, 10 mM 3nY, and 2 μg mL^-1^ ceftazidime, and grown at 37°C for 3 hours, and then made electrocompetent with serial washes of 10% glycerol. Samples were transformed with GFP variants, recovered with LB media for one hour at 37°C, then plated on LB-agar supplemented with 10 mM 3nY, .02% glucose, 50 μg mL^-1^ KAN, and 2 or 22 μg mL^-1^ CAZ, and incubated overnight at 37°C. Four samples from each plate were picked and grown overnight at 37°C in 500 μL LB with 10 mM 3nY, .02% glucose, 50 μg mL^-1^ KAN, and 2 or 22 μg mL^-1^ CAZ. A 2 μL aliquot of each overnight culture was used to inoculate 1 mL LB media supplemented with 50 μg mL^-1^ KAN, and containing either 1 mM 3nY, 1 mM 3iY, or with no supplemented ncAA. Cultures were grown at 37°C for 3 hours, then induced with 1 mM IPTG, and grown for another 5 hours at 37°C. Final cultures were rinsed twice with iterative centrifugation at 2000*g* and PBS (50 mM phosphate, 300 mM NaCl, pH 8) at 4°C. Samples were resuspended in 1 mL PBS and a 150 μL aliquot was used to measure relative fluorescence by measuring absorbance at 600 nm (OD600), and fluorescence using a 475 nm excitation and measuring emission at 525 nm using Tecan Infinite M200 pro microplate reader.

## Acknowledgements

Acknowledgements. We would like to thank the Genome Sequencing and Analysis Facility (GSAF) at the University of Texas at Austin for library preparation and sequencing of bacterial genomes. We thank Dr. Jeffery Barrick for his thoughtful discussions and insights, and for providing the Keio knockout strains used in this study. We would also like to thank Terry Stewart Jr. for diligently preparing laboratory materials. This work was supported by a National Security Science and Engineering Faculty Fellowship (FA9550-10-1-0169), the Air Force Office of Science research (FA9550-14-1-0089), and Defense Advanced Research Projects Agency (HR0011-15-C0095). ADE is supported by the Welch Foundation (F-1654).

